# Integration of sound and locomotion information by auditory cortical neuronal ensembles

**DOI:** 10.1101/2022.05.16.492071

**Authors:** Carlos Arturo Vivaldo, Joonyeup Lee, MaryClaire Shorkey, Ajay Keerthy, Gideon Rothschild

**Affiliations:** Department of Psychology, University of Michigan, Ann Arbor, MI 48109, USA; Neuroscience Graduate Program, University of Michigan, Ann Arbor, MI 48109, USA; Kresge Hearing Research Institute and Department of Otolaryngology - Head and Neck Surgery, University of Michigan, Ann Arbor, MI 48109, USA

## Abstract

The ability to process and act upon incoming sounds during locomotion is critical for survival. Intriguingly, sound responses of auditory cortical neurons are on average weaker during locomotion as compared to immobility and these results have been suggested to reflect a computational resource allocation shift from auditory to visual processing. However, the evolutionary benefit of this hypothesis remains unclear. In particular, whether weaker sound-evoked responses during locomotion indeed reflect a reduced involvement of the auditory cortex, or whether they result from an alternative neural computation in this state remains unresolved. To address this question, we first used neural inactivation in behaving mice and found that the auditory cortex plays a critical role in sound-guided behavior during locomotion. To investigate the nature of this processing, we used two-photon calcium imaging of local excitatory auditory cortical neural populations in awake mice. We found that underlying a net inhibitory effect of locomotion on sound-evoked response magnitude, spatially intermingled neuronal subpopulations were differentially influenced by locomotion. Further, the net inhibitory effect of locomotion on sound-evoked responses was strongly shaped by elevated ongoing activity. Importantly, rather than reflecting enhanced “noise”, this ongoing activity reliably encoded the animal’s locomotion speed. Prediction analyses revealed that sound, locomotive state and their integration are strongly encoded by auditory cortical ensemble activity. Finally, we found consistent patterns of locomotion-sound integration in electrophysiologically recorded activity in freely moving rats. Together, our data suggest that auditory cortical ensembles are not simply suppressed by locomotion but rather encode it alongside sound information to support sound perception during locomotion.

## Introduction

Continuous processing of incoming sensory information is critical for survival and adaptive behavior. Whereas studies of the neural mechanisms of sensory processing have traditionally focused on immobile subjects, some of the most critical behaviors in humans and other animal species -- such as foraging for food, seeking a mate, and evading danger -- occur during locomotion. To gain a coherent perception of the environment during locomotion and be able to rapidly guide appropriate behavior, the brain must integrate incoming external cues with one’s own motion. For example, humans integrate incoming sounds with locomotion during simple walking, as manifested by the influence of modified auditory feedback on walking pace (Cuppone et al., 2018; Redd and Bamberg, 2012; Tajadura-Jiménez et al., 2015; Turchet et al., 2015; Turchet et al., 2018; Turchet et al., 2013). Moreover, auditory feedback has been shown to improve walking in aged patients and those with neurodegenerative disorders (Cornwell et al., 2020; Rodger et al., 2014; Schauer and Mauritz, 2003). Integration of sounds with self-motion has also been studied in the context of other behaviors such as dance (Karpati et al., 2015; Ravignani and Cooke, 2016) and sound-guided finger-tapping (Carr et al., 2016; Chen et al., 2008; Tierney and Kraus, 2013, 2016). In non-humans, perhaps the best known example is bat echolocation (Falk et al., 2014; Ghose et al., 2006; Moss and Surlykke, 2001), yet various forms of audiomotor integration have been studied in diverse animal species, including Praying mantids (Triblehorn and Yager, 2005), Dholes (Fox, 1984) and mice (Whitton et al., 2014). Thus, the ability to process incoming sounds during locomotion and integrate them with the locomotive state of the body is fundamental in both humans and other animal species.

The auditory cortex is a key candidate brain region for processing incoming sounds during locomotion due to its well-established role in context-, behavior-, and decision-making-dependent sound processing (Cohen et al., 2011; Jaramillo and Zador, 2011; Kuchibhotla et al., 2017; Rodgers and DeWeese, 2014; Ulanovsky et al., 2003; Xiong et al., 2015b; Znamenskiy and Zador, 2013a). Intriguingly, previous studies have found that locomotion has a generally suppressive effect on sound-evoked responses in the auditory cortex (Bigelow et al., 2019; Schneider et al., 2014; Zhou et al., 2014). Based on these results, and the finding that responses in the primary visual cortex are generally enhanced during locomotion (Dadarlat and Stryker, 2017; Niell and Stryker, 2010; Vinck et al., 2015), it has been suggested that locomotion reflects a resource allocation shift from audition to vision (Schneider et al., 2014; Zhou et al., 2014). However, the evolutionary benefit of this hypothesis remains debated (Bigelow et al., 2019), especially in nocturnal animals with poor vision such as rodents. In particular, whether weaker sound-evoked responses during locomotion indeed reflect a reduced involvement of the auditory cortex, or whether they result from an alternative neural computation in this state remains unresolved. In support of the latter possibility, studies in the visual and somatosensory cortices have revealed that locomotion itself is strongly encoded within these regions and integrated with the respective sensory cues (Ayaz et al., 2013; Ayaz et al., 2019; Saleem et al., 2013). Here we tested the hypothesis that auditory cortical ensembles are not simply suppressed by locomotion but rather explicitly encode it and incorporate it with sound information into an integrated neural code.

As a first step to test this hypothesis, we used neural inactivation in behaving mice to determine the involvement of the auditory cortex in sound-guided behavior during different locomotive states and found that the auditory cortex plays a critical role in sound processing during locomotion. To investigate the nature of this processing, we used two-photon calcium imaging in the auditory cortex of mice and quantified encoding of sound, locomotive state and their integration by local neuronal populations. To test whether these findings generalize to freely moving animals we examined these questions in electrophysiologically recorded auditory cortical ensembles of freely moving rats. Together, our data suggest that auditory cortical ensembles explicitly and reliably encode self-locomotion and integrate it with sound information to support sound perception during locomotion.

## Results

### Auditory cortical activity is required for sound processing during locomotion

To determine the role of the auditory cortex (AC) in sound processing during locomotion, we tested the influence of AC inactivation on sound-guided behavior during locomotion and immobility. Specifically, male and female mice were first implanted with bilateral cannula into the AC for subsequent drug delivery and allowed to recover for at least 5 days. Mice were put on water restriction and were trained on an appetitive trace conditioning task during head fixation while standing on a rotatable plate that allowed the animals to stand or run at will. In each training trial, the presentation of an 8 kHz tone was followed by a drop of water reward, delivered 1 s after sound termination. Using a closed-loop system that received the output of a rotary encoder at the base of the plate, one group of mice was trained on this task wherein sounds were presented only during immobility (n=7) and the other group was trained on a similar version of the task in which sounds were presented during locomotion (n=8). Mice of both groups were trained until they learned the sound-reward association as evidenced by an increase in lick rate following the sound and before reward delivery (“predictive licking”, **Fig. 1A, B**). To test whether AC activity is necessary for this sound-guided reward predictive behavior, we measured the influence of AC inactivation on predictive licking. To this end, we measured behavioral performance in trained mice following infusion of the GABAA receptor agonist muscimol (MUS), or inert phosphate buffer solution (PBS) as a control, into the AC (**Fig. 1C**). We found that inactivation of the AC induced a significant reduction in sound-triggered predictive licking when sounds were presented in locomotion but not in immobility (**Fig. 1C,D**). Furthermore, the reduction in predictive licking following AC inactivation was significantly larger in locomotion as compared to in immobility (**Fig. 1E**). These findings suggest that the AC plays an important role in sound-guided behavior during locomotion in mice. The finding that the AC is required for sound-guided behavior during locomotion even more than in immobility suggests that previous reports of weaker average sound-evoked responses may not reflect a reduced involvement of AC in auditory perception but rather that it carries out a different form of computation.

**Fig 1.**
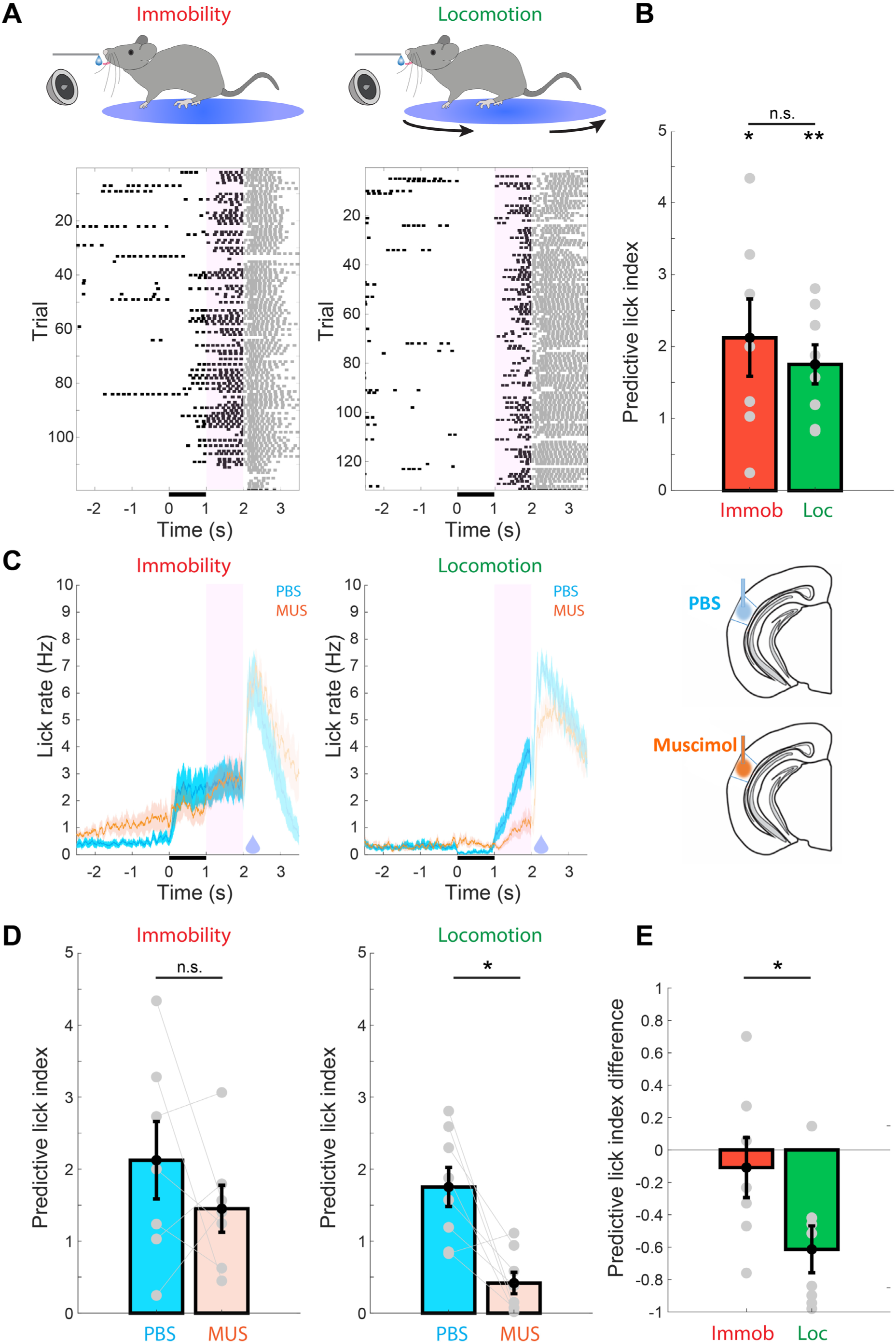
Auditory cortical activity is necessary for sound processing during locomotion. **(A)** Top: Illustration of the behavioral setup for sound-guided predictive licking in immobility (left) and locomotion (right). Bottom: Peri-sound lick histograms of example behavioral sessions from trained animals performing the task in immobility (left) and locomotion (right). Licking in the pink shaded area following sound termination represents prediction of upcoming reward (delivered at 2 s). Licks following reward delivery are shaded as they do not require sound processing or reward prediction **(B)** Animals performing the task in both immobility (n=7 mice) and locomotion (n=8 mice) exhibited significant predictive licking (Immobility: P=0.0156, Locomotion: P=0.0078, two-sided signed rank test of difference from 0) and the predictive lick index did not differ between these groups (P=0.6126, rank sum test). **(C)** Peri-sound lick histograms across animals performing the task in immobility (left) and locomotion (right) when the AC was infused with either PBS or muscimol. Solid lines denote the mean and the shaded area represents s.e.m across animals. Predictive licking in locomotion but not in immobility is clearly reduced following AC inactivation using muscimol. **(D)** Predictive lick index in immobility and locomotion following infusion of PBS or muscimol. There was a significant reduction in predictive lick index following infusion of PBS/MUS in locomotion (P=0.0156, signed rank test) but not in immobility (P=0.578, signed rank test). Error bars represent mean±s.e.m across animals. Lines connecting gray circles represent data from the same animal in the different conditions. **(E)** The per-animal differences in predictive lick index following infusion of muscimol and PBS (negative values denote a reduction in predictive licking following muscimol relative to PBS). The reduction in predictive lick index following AC inactivation with muscimol was significantly larger in locomotion than in immobility (P=0.040, rank sum test).

### Locomotion differentially modulates sound-evoked responses of spatially intermingled subnetworks of excitatory L2/3 neurons in the auditory cortex

To study the nature of information processing by local groups of L2/3 excitatory neurons (“neuronal ensembles”) of the auditory cortex (AC) during locomotion, we carried out two-photon calcium imaging in head-fixed Thy1-GCaMP6f mice that were free to stand or run at will on a rotatable plate (**Fig. 2A-C**). We first examined how locomotion modulates the responses of neurons to broad-band noise (BBN) bursts in 985 AC neurons from 7 mice, of which 612 neurons had a sufficient number of responses in both immobility and locomotion to allow for comparison. In keeping with most previous studies, we started with examining baseline-subtracted responses, which are defined as the difference between the activity evoked by the sound and the activity immediately preceding the sound. Locomotion had a diverse influence on sound-evoked responses of individual neurons, including invariance (**Fig. 2D** neurons 1+2), suppression (**Fig. 2D** neurons 3+4) and enhancement (**Fig. 2D** neurons 5+6), consistent with a recent report (Bigelow et al., 2019). Across all neurons that exhibited significant BBN-evoked responses in immobility (194/612, 31.7%), the population-average responses were significantly reduced during locomotion (**Fig. 2E**). To test whether these findings were unique to responses to BBN, we examined how locomotion modulates responses to pure tones and complex sounds. These experiments revealed a similar influence of locomotion on sound evoked responses, namely a net population-average decrease that coexists with heterogeneous influences at the single-cell level (**Suppl. Fig. 1**).

**Fig 2.**
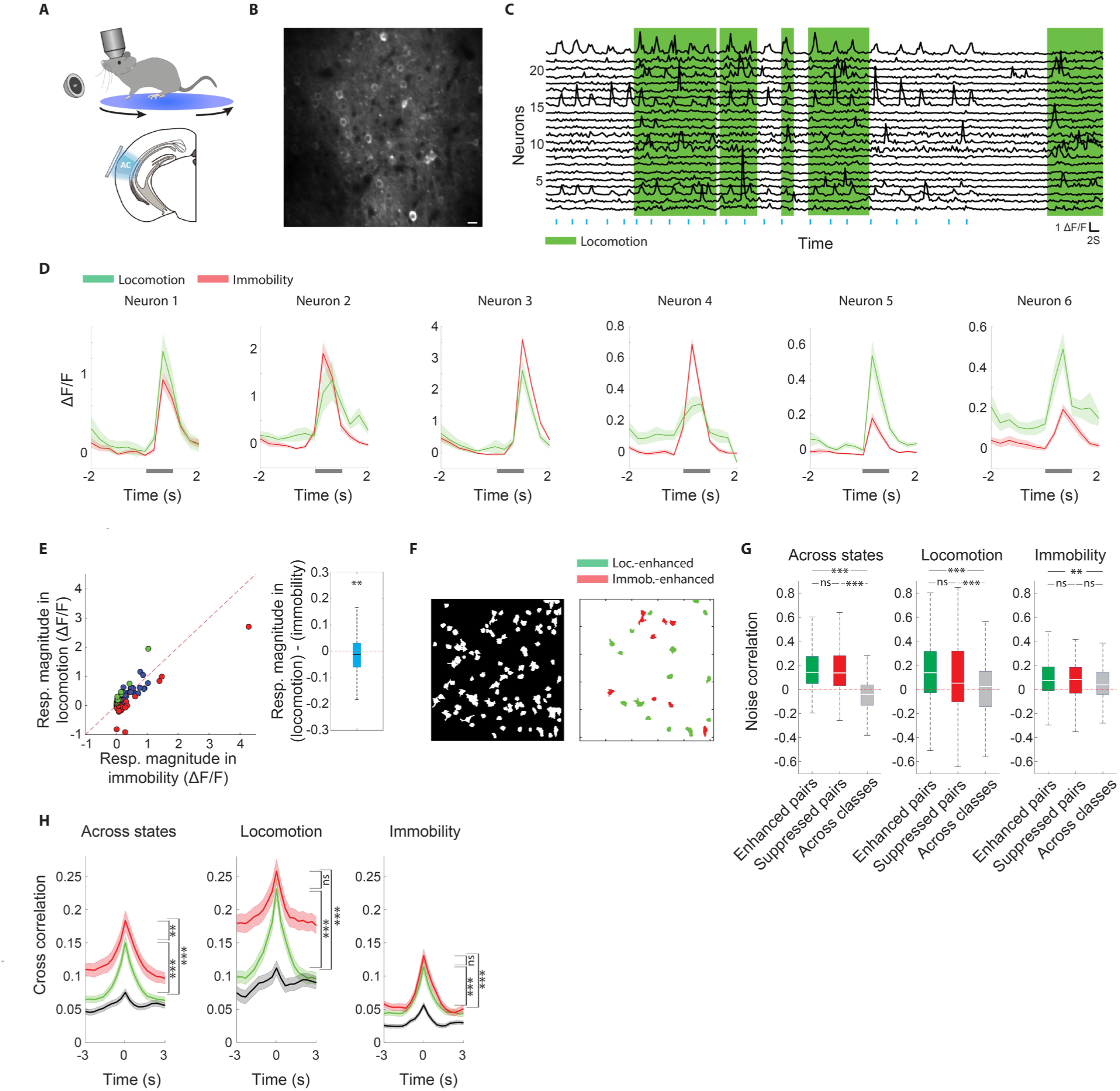
Locomotion differentially modulates sound-evoked responses of spatially intermingled subnetworks of excitatory L2/3 neurons in the auditory cortex. **(A)** Illustration of the experimental setup **(B)** Two-photon average micrograph of an example local neuronal ensemble in L2/3 of the auditory cortex. Scale bar: 10µm. **(C)** Relative change in fluorescence (ΔF/F) of 22 neurons from the micrograph in ‘B’ during an imaging session. Periods of locomotion are marked in green. **(D)** Sound-triggered peri-stimulus time histograms from 6 example neurons. Sound presentation trials in which the animal was immobile (red) and running (green) were grouped separately. Locomotion had diverse effects on sound-evoked responses of different neurons, including invariance (neurons 1+2), reduction (neurons 3+4) and enhancement (neurons 5+6) **(E)** Left: Sound-evoked responses in immobility and locomotion across all BBN-responsive neurons. Red and green dots represent neurons that individually exhibited a significantly stronger and weaker response during immobility, respectively.Blue dots represent neurons that did not exhibit a significant difference. Right: Box plot describing sound-evoked responses in locomotion minus immobility across all BBN-responsive neurons. The distribution was significantly lower than 0 (P=0.009, two-sided Wilcoxon signed-rank). For this and subsequent whisker plots, the central mark indicates the median, the bottom and top edges of the box indicate the 25th and 75th percentiles, respectively and the whiskers extend to the most extreme data points not considered outliers. **(F)** Left: shadow illustration showing the identified cell bodies from an example imaging session. Right: A corresponding illustration where neurons marked in red and green exhibit significantly weaker and stronger sound-evoked responses during locomotion, respectively. Neurons that show no significant difference are not shown. **(G)** Noise correlation between simultaneously imaged pairs of neurons whose sound-evoked responses were both enhanced (green), suppressed (red), or mixed (one neuron enhanced and the other suppressed, gray). All tested with One-way ANOVA and Tukey-Kramer post hoc test. Across states: F(2,627)=129.43, P=8.8e-48, enhanced vs. across: P=9.6e-10, suppressed vs. across: P=9.6e-10, enhanced vs. suppressed: P=0.667. Locomotion: F(2,627)=29.41, P=6.2e-13, enhanced vs. across: P=9.6e-10, suppressed vs. across: P=4.6e-4, enhanced vs. suppressed: P=0.073. Immobility: F(2,627)=5.54, P=0.0041, enhanced vs. across: P=0.0027, suppressed vs. across: P=0.277, enhanced vs. suppressed: P=0.533. **(H)** Cross-correlograms of the continuous ΔF/F traces between pairs of neurons of different locomotion-modulation categories. Across states: F(2,627)=32.22, P=4.8e-14, enhanced vs. across: P=6e-8, suppressed vs. across: P=6e-8, enhanced vs. suppressed: P=0.0063; Locomotion: F(2,627)=25.83, P=1.7e-11, enhanced vs. across: P=2.4e-7, suppressed vs. across: P=6e-8, enhanced vs. suppressed: P=0.0737; Immobility: F(2,627)=25.71, P=1.9e-11, enhanced vs. across: P=6e-8, suppressed vs. across: P=6e-8, enhanced vs. suppressed: P=0.2815

As information is processed in the brain by groups of interacting neurons, we examined how locomotion modulates responses at the ensemble level. Notably, we observed that within the same local ensemble individual neurons often exhibited opposing locomotion-related modulation of sound-evoked responses (**Fig. 2F**). We thus wondered whether neurons whose BBN-evoked responses are similarly modulated by locomotion (suppressed/enhanced) form functional subnetworks. To this end we calculated trial-by-trial noise correlations in sound-evoked responses between pairs of simultaneously imaged neurons that both exhibited significantly suppressed responses during locomotion (114 pairs), pairs of neurons that both exhibited significantly enhanced responses during locomotion (244 pairs), and pairs of neurons in which one neuron exhibited significantly enhanced responses and the other significantly suppressed responses during locomotion (272 pairs). When calculated across all sound presentations in both behavioral states (immobility and locomotion), we confirmed that pairs of neurons whose BBN-evoked responses are similarly modulated by locomotion exhibited significantly higher noise correlations than pairs of neurons with opposing locomotion-modulation (**Fig. 2G**, left). Interestingly, we found that this pattern also held true when examining responses during locomotion only (**Fig. 2G**, middle). When examining responses during immobility only, pairs of locomotion-enhanced neurons showed higher noise correlations than pairs of neurons across classes (**Fig. 2G**, right). None of these patterns were observed in trial-shuffled data (**Suppl. Fig. 2**). Further, enhanced synchronization between pairs of neurons whose BBN-evoked responses are similarly modulated by locomotion was also observed when examining the full continuous activity traces, as evidenced by significantly enhanced peaks in activity cross-correlograms (**Fig. 2H**). Thus, beyond the influence on individual neurons, locomotion has a synchronizing effect on pairs of neurons with shared locomotion-induced modulation. These data suggest that locomotion differentially modulates sound-evoked responses of spatially-intermingled subnetworks of AC neurons.

### Enhanced ongoing activity during locomotion contributes to a reduction of baseline-subtracted sound-evoked responses

Despite the local functional heterogeneity in the influence of locomotion on sound-evoked responses, the net effect of locomotion was a reduction in baseline-subtracted responses (**Fig. 2E**), consistent with previous reports (Bigelow et al., 2019; Schneider et al., 2014). We sought to further investigate the source of this reduction and noticed that many neurons exhibited increased ongoing activity during locomotion, which manifested as increased activity before stimulus onset (**Fig. 2D**). Across the population of BBN-responsive neurons, the average sound-triggered peri-stimulus time histogram (PSTH) exhibited a clear elevation in pre-stimulus activity during locomotion as compared to immobility (**Fig. 3A**, orange arrow). Indeed, ongoing activity was significantly higher during locomotion as compared to immobility across responsive neurons, as well as across all neurons (**Fig. 3B**). While running on the rotating plate seemed to generate no noticeable sound, we wondered whether increased ongoing activity during locomotion can be attributed to processing of self-generated sounds. To this end we imaged ongoing activity of 269 neurons from 5 animals in the presence of sound-masking continuous background broad-band noise. We found that locomotion also had a population-wide average excitatory effect in the presence of background masking noise, in BBN-responsive neurons (73/269, 27% **Fig. 3C**, left) as well as across all neurons (**Fig. 3C**, right). Thus, locomotion-related increase in ongoing activity persists in the presence of background masking noise, suggesting that it is at least partly independent of self-generated sounds.

**Fig 3.**
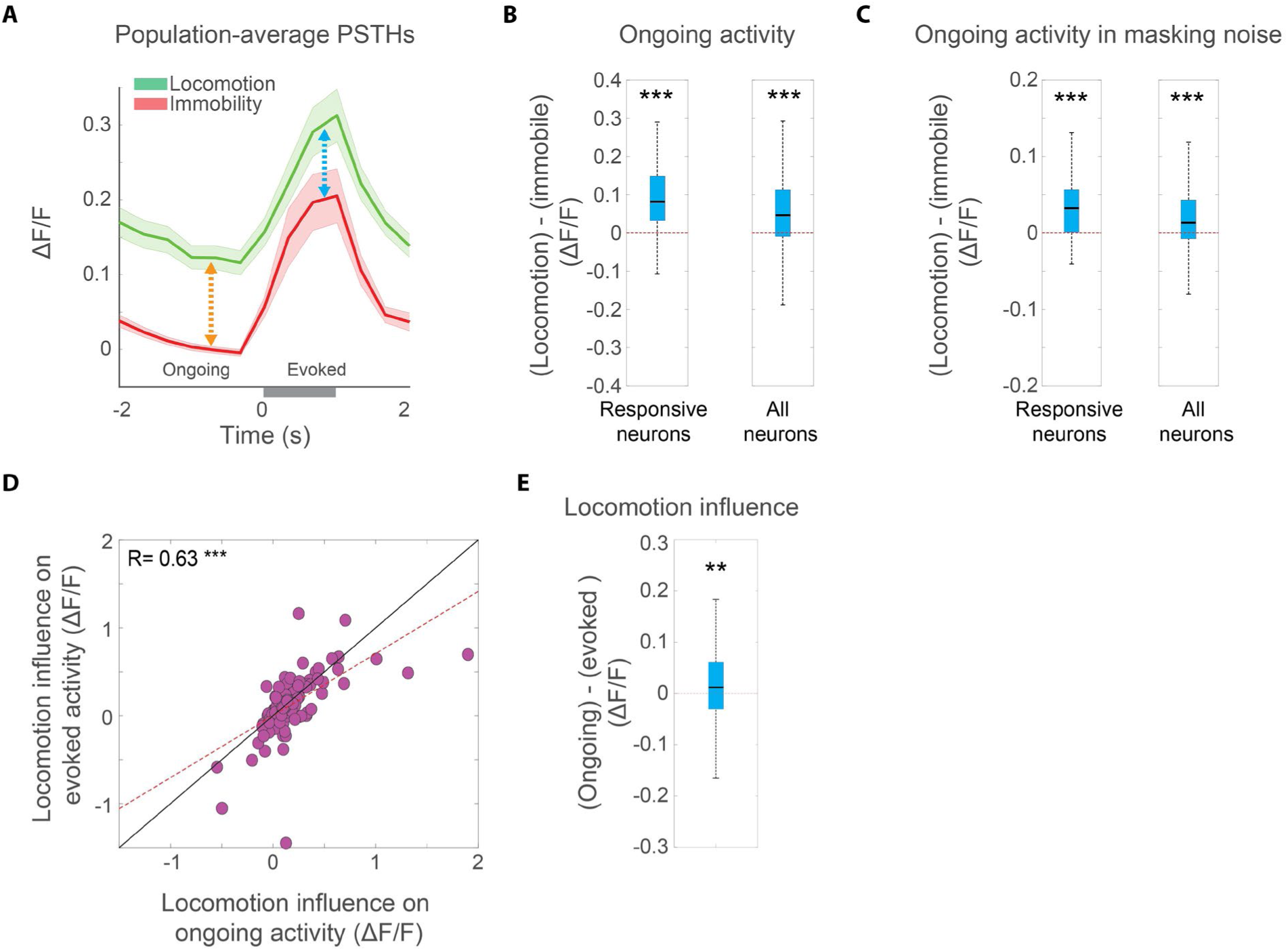
Enhanced ongoing activity during locomotion contributes to a reduction of baseline-subtracted sound-evoked responses. **(A)** Population-level peri-stimulus time histogram across all BBN-responsive neurons during immobility (red) and locomotion (green). Solid lines and shaded areas indicate mean±SEM. **(B)** The per-neuron difference between ongoing (pre-stimulus) activity during locomotion and immobility was significantly higher than 0 for both BBN-responsive neurons (left, P= 2.9e^-23^, two-sided Wilcoxon signed-rank test) and all neurons (right, P= 7e^-27^, two-sided Wilcoxon signed-rank test). For this and subsequent whisker plots, the central mark indicates the median, the bottom and top edges of the box indicate the 25th and 75th percentiles, respectively and the whiskers extend to the most extreme data points not considered outliers. **(C)** The per neuron difference between ongoing activity during locomotion and immobility in the presence of masking background noise is significantly higher than 0 for both BBN-responsive neurons (left, P=4.7e^-7^, two-sided Wilcoxon signed-rank test) and all neurons (right, P=7.7e^-11^, two-sided Wilcoxon signed-rank test). **(D)** Locomotion-related influence on ongoing activity (in the pre-stimulus time window, orange arrow in Fig. 2A) against locomotion-related influence on sound response activity (blue arrow in Fig. 2A) for all BBN-responsive neurons. Dashed red line indicates best linear fit. Pearson correlation coefficient: R=0.63, P=6.7e^-23^. Black line indicates the diagonal. **(E)** The per-neuron influence of locomotion on ongoing activity (orange arrow in Fig. 2A) minus the influence of locomotion on sound-evoked activity (blue arrow in Fig. 2A) was significantly positive across all BBN-responsive neurons (P=0.0095, two-sided Wilcoxon signed-rank test).

As ongoing, pre-stimulus activity is subtracted out in the standard calculation of sound-evoked responses, an increase in pre-stimulus activity could contribute to reduced sound-evoked responses during locomotion. To further test this possibility, we compared the influence of locomotion on activity during the pre-stimulus time window (**Fig. 3A**, orange arrow, “ongoing activity”) and stimulus time window (**Fig. 3A**, blue arrow, “evoked activity”). We found that across all BBN-responsive neurons, the influence of locomotion on activity during the pre-stimulus and stimulus time windows were highly correlated (**Fig. 3D**). Importantly, however, while locomotion was associated with increased activity in both the pre-stimulus and stimulus time windows, the locomotion-related increase in activity was significantly higher during the pre-stimulus time window (**Fig. 3D,E**). These data demonstrate that the observed average reduction in baseline-subtracted sound responses during locomotion is at least partly shaped by increased ongoing, pre-sound activity. We therefore wondered whether this enhanced baseline activity during locomotion, which is traditionally subtracted out in the calculation of sound response magnitude and seems to impair neural sound detection, may in fact reflect encoding of meaningful information for auditory cortical processing.

### Auditory cortical L2/3 neurons and ensembles reliably encode locomotion speed

We wondered whether the enhanced baseline activity during locomotion, that seems to impair neural sound detection, may in fact reflect encoding of meaningful information for auditory cortical processing. In particular, we hypothesized that enhanced ongoing activity during locomotion encodes the animal’s locomotion velocity. To test this hypothesis, we first asked whether neural activity of individual neurons is significantly correlated with locomotion speed. We calculated the correlations between the continuous relative change in fluorescence of each neuron and the running speed of the mouse, utilizing a large subset of our imaged neurons (647/985) that were imaged while the continuous running speed of the animal was acquired. We found that activity of auditory cortical neurons could exhibit surprisingly high positive correlations with locomotion speed (**Fig. 4A**), and in fewer cases significant negative correlations with locomotion speed (**Fig. 4B**). Across the population, 52% of neurons (335/647) showed significant positive correlation with locomotion speed, 24% of neurons (155/647) exhibited significant negative correlation with locomotion speed and 24% (157/647) showed no significant correlation with locomotion speed (**Fig. 4C**). The distribution of correlations between neural activity and locomotion speed was skewed to the right (**Fig. 4D**, skewness=0.84), consistent with our finding of a population-level enhancement in baseline activity during locomotion.

**Fig 4.**
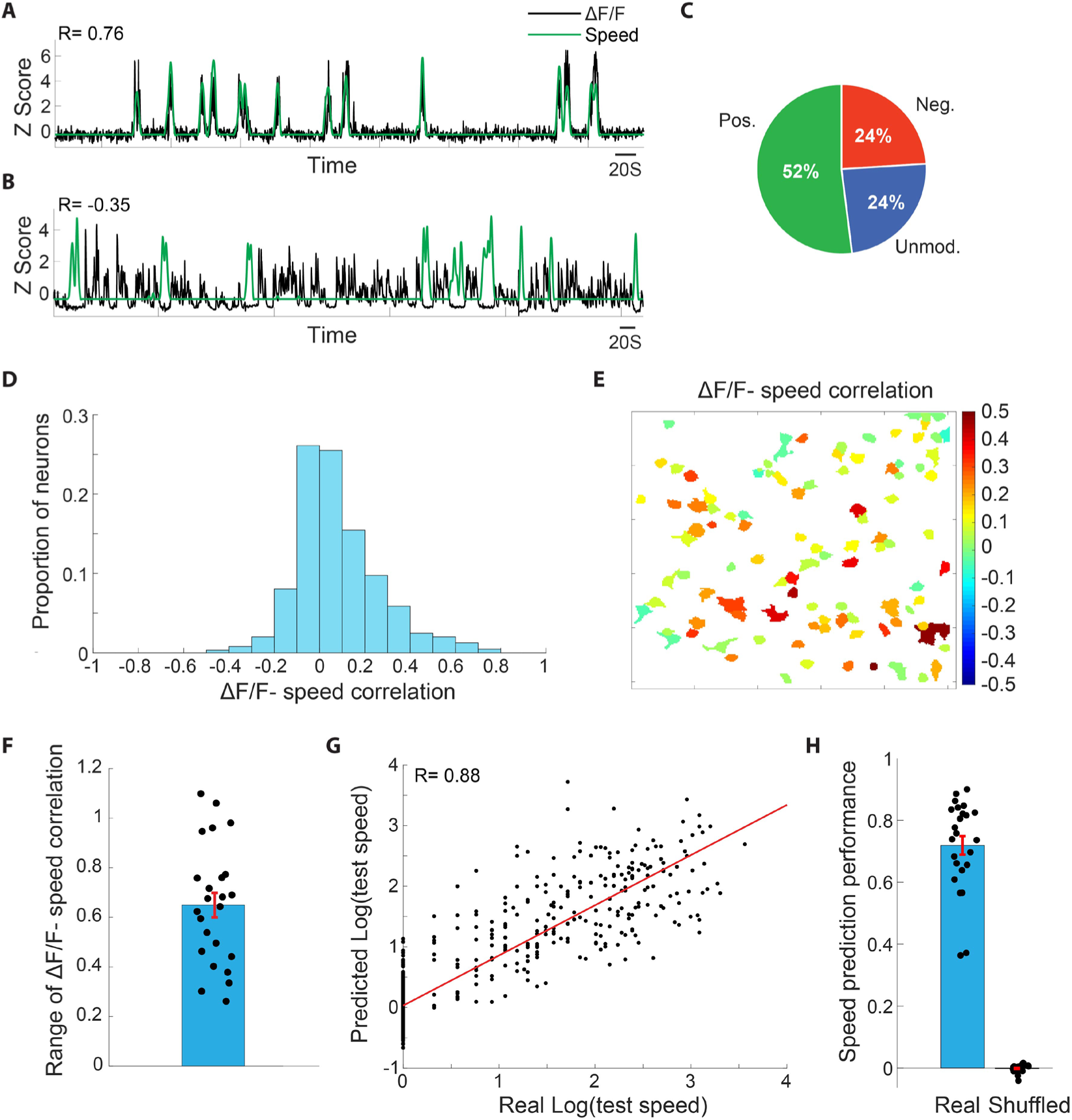
Auditory cortical L2/3 neurons and ensembles reliably encode locomotion speed. **(A)** Z-scored ΔF/F of an example neuron (black trace) overlaid on the Z-scored locomotion speed of the mouse (green trace) during an example imaging session. This neuron exhibited a correlation of R=0.76 with locomotion speed across the session. **(B)** An example from a different neuron, showing a negative correlation with locomotion speed of R=-0.35. **(C)** Proportions of AC L2/3 neurons showing significant positive, significant negative and non-significant correlation with locomotion speed **(D)** The distribution of ΔF/F-locomotion speed correlations across the population **(E)** An illustration of all neurons in an example imaging session (same as in Fig. 1F), color coded according to each neuron’s ΔF/F-locomotion speed correlation value. Local ensembles exhibited a high degree of heterogeneity in correlation with locomotion speed. **(F)** The ensemble-level range in ΔF/F-locomotion speed correlation values across ensembles. **(G)** The predicted log(speeds) of an example test-set against the real log(speeds) of that test-set, showing a correlation of 0.88. **(H)** Speed prediction performance, measured as the correlation values between the predicted and real locomotion speeds across ensembles. Shuffled values were derived by randomly shuffling the predicted speed values.

Interestingly, we also found that within local excitatory auditory cortical ensembles in L2/3, individual neurons exhibited high diversity in correlations between neural activity and locomotion speed (**Fig. 4E**). Within local neuronal ensembles, the average range of correlations between locomotion speed and relative change in fluorescence of the different neurons was 0.65 (**Fig. 4F**). These findings suggest that despite the net excitatory effect, locomotion modulates ongoing activity of local excitatory neuronal populations in a spatially fine-tuned manner rather than acting as a global uniform modulator.

To further quantify the degree of information that auditory cortical ensembles convey about locomotion speed, we implemented a cross-validated generalized linear model (GLM) to test if locomotion speed can be decoded from ongoing ensemble activity. For each imaging session of a single neuronal ensemble, a GLM was trained on a random half of the imaging session data and tested on the other half, and this procedure was repeated 200 times for robust estimation. In the test phase, the GLM model that was constructed in the training phase predicted locomotion speed based on ensemble patterns of neural activity of the test set. We found that in many cases the predictions of the model were highly correlated with the actual speeds (**Fig. 4G**). Across ensembles, the correlation between the predicted speed and real speed averaged 0.72 (**Fig. 4H**). These findings suggest that ongoing locomotion speed is reliably encoded by the activity of local neuronal ensembles in the auditory cortex.

### Integration of sound and locomotion information by excitatory neuronal ensemble in L2/3 of the auditory cortex

Taken together, our results suggest that excitatory neurons in L2/3 of the auditory cortex robustly encode both external sounds and locomotion. These findings raise the question of whether these two variables-external sounds and locomotion state-are simultaneously represented and integrated within the local network level. As our previous results suggest, neurons within the same ensemble exhibited a range of modulations by both locomotion state and sound (**Fig. 5A**). We thus hypothesized that ensemble-level activity patterns could provide discriminability about both of these attributes. To test this hypothesis we quantified the amount of information that each ensemble encoded about both locomotion state (immobility/locomotion) and about sound occurrence during locomotion (n=19 ensembles). To this end we implemented cross-validated support vector machine (SVM) analyses on each ensemble’s neural activity patterns and quantified the predictive power that it provided to discriminate between immobility and locomotion and between sound occurrence and no sound during locomotion (**Fig. 5B**). We found that across ensembles, both locomotive state and sound occurrence during locomotion could be significantly decoded from ensemble activity. Furthermore, the activity patterns of all ensembles (19/19) could individually significantly discriminate locomotive state, and activity patterns of most ensembles (12/19) could individually significantly discriminate both locomotive state and sound occurrence (**Fig. 5C**, P<0.05 bootstrap analysis). As an additional test for the ability of ensembles to encode both locomotive state and sound, we asked whether ensembles could discriminate between the four combinations of these variables: no sound in immobility, sound in immobility, no sound in locomotion and sound in locomotion. We found that AC ensembles could significantly discriminate between these 4 options better than chance and 18/19 ensembles showed individually significant discrimination, with some ensembles discriminating at 70-80% correct rates (**Fig. 5D**, chance=25%). These data suggest that neuronal ensembles in L2/3 of the auditory cortex co-encode and integrate incoming sound with locomotion information.

**Fig 5.**
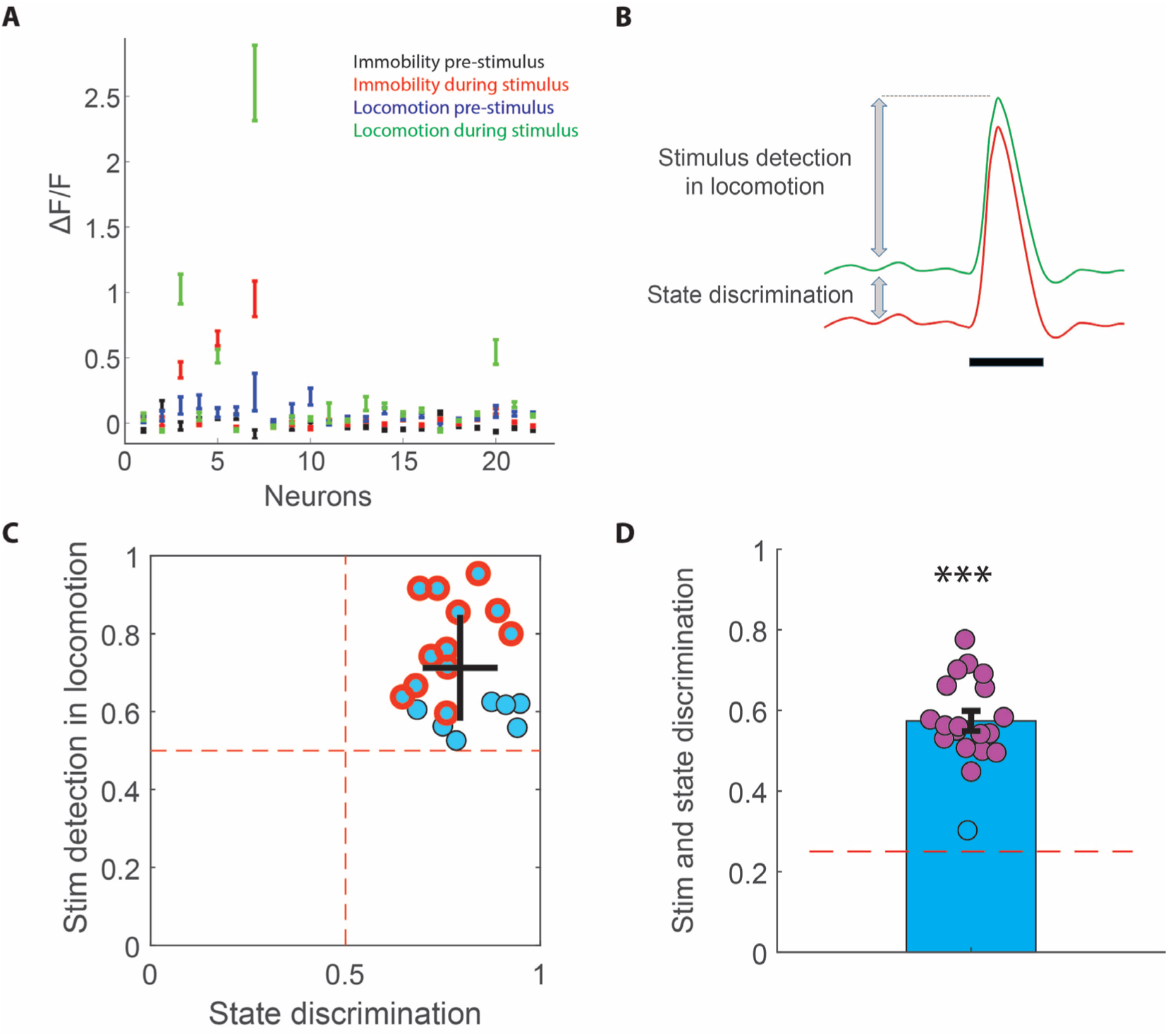
Integration of sound and locomotion information by excitatory neuronal ensemble in L2/3 of the auditory cortex. **(A)** Activity levels (mean±SEM) of individual neurons in an example ensemble in four different conditions: pre-stimulus (ongoing) activity in immobility (black), sound-evoked activity during immobility (red), pre-stimulus (ongoing) activity in locomotion (blue), sound-evoked activity during locomotion (green). **(B)** Schematic illustration of the measures used for stimulus detection in locomotion and state discrimination. **(C)** Performance of stimulus detection in locomotion against state discrimination across ensembles. Blue points indicate ensembles showing significant state discrimination, red-circled points indicate ensembles showing significant stimulus detection in locomotion. Black cross shows mean±STD of the two measures. Across ensembles, both locomotive state (P=1.318e^-4^) and sound detection during locomotion (P=1.316e^-4^) could be significantly decoded from ensemble activity (Two-sided Wilcoxon signed-rank test of difference from chance of 0.5). 10/19 ensembles individually significantly discriminated both locomotive state and sound occurrence in locomotion (P<0.05, bootstrap analysis). **(D)** Significant discrimination of the four sound/locomotion state combinations (pre-sound in immobility, sound in immobility, pre-sound in locomotion, sound in locomotion) by ensemble activity patterns (P=1.318e-4, two-sided Wilcoxon signed-rank test of difference from 0.25). 18/19 could individually significantly discriminate between these 4 options (P<0.05, bootstrap analysis)

### Integration of sound and locomotion in the freely moving rat

Finally, we wished to test whether our findings of sound-locomotion integration in head-fixed animals generalize to freely-moving animals. To this end, we analyzed electrophysiological recordings from freely-moving rats that were implanted with tetrodes in the auditory cortex (Rothschild et al., 2017). Recordings were carried out as rats traversed a Y-shaped track for food reward delivered at reward wells (**Fig. 6A**). In a pseudorandom ∼25% of trials, following nose-poking in the Home well rats were presented with series of chirp-pair sounds, which signaled that subsequent reward is delivered in the Sound well. We identified putative excitatory and inhibitory interneurons based on spike waveform (**Suppl. Fig. 3**) and focused all analyses on putative excitatory neurons. We recorded a total of 248 putative excitatory neurons that had a sufficient number of responses in both immobility and locomotion to allow comparison. Of these, 21% (51/248) were significantly responsive to the target sound during immobility.

**Fig 6.**
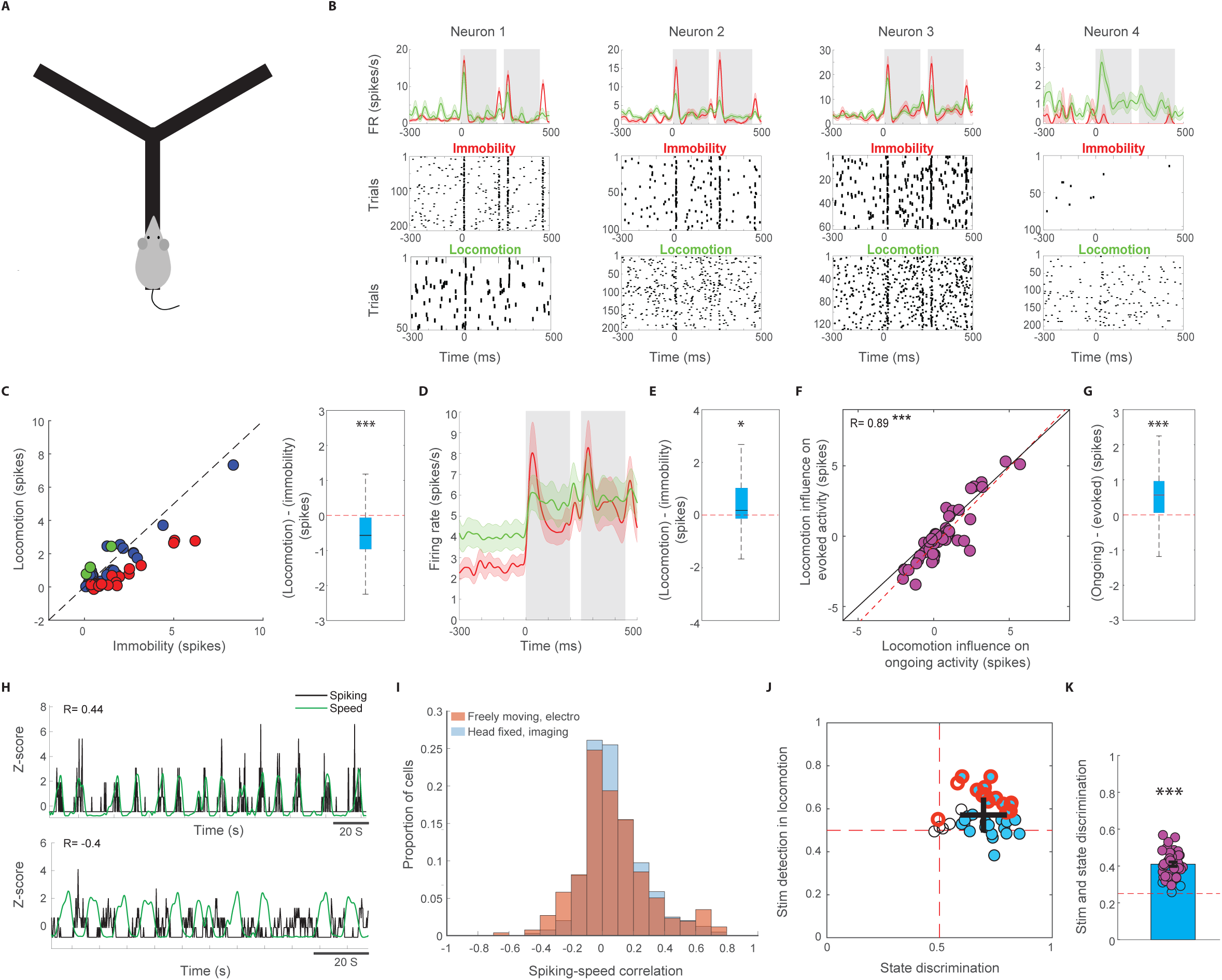
Integration of sound and locomotion in the freely moving rat **(A)** Illustration of the experimental setup for electrophysiological recordings in freely-moving rats **(B)** Sound-triggered peri-stimulus time histograms from 4 example neurons. Sound presentation trials in which the animal was immobile (red) and running (green) were grouped separately. Neurons showed diverse patterns of modulation of sound-evoked responses during locomotion **(C)** Left: Sound-evoked responses in immobility and locomotion across all target-sound responsive neurons. Red and green circles denote neurons that individually exhibited a significantly stronger and weaker response during immobility, respectively. Blue circles denote neurons that did not exhibit a significant difference. Right: The per-neuron difference in sound-evoked response between locomotion and immobility across all responsive neurons was significantly lower than 0 (P=3.4e-5, two-sided Wilcoxon signed-rank). For this and subsequent whisker plots, the central mark indicates the median, the bottom and top edges of the box indicate the 25th and 75th percentiles, respectively and the whiskers extend to the most extreme data points not considered outliers. **(D)** Population-level peri-stimulus time histogram across all target-sound responsive neurons during immobility (red) and locomotion (green). Solid lines and shaded areas indicate mean±SEM. **(E)** The per-neuron difference between ongoing (pre-stimulus) activity during locomotion and immobility was significantly higher than 0 across target-sound responsive neurons (P=0.014, two-sided Wilcoxon signed-rank test) **(F)** Locomotion-related influence on ongoing activity (in the pre-stimulus time window) against locomotion-related influence on sound response activity for all target-sound responsive neurons. Dashed red line indicates best linear fit. Pearson correlation coefficient: R=0.89, P=1.8e-18. Black line indicates the diagonal. **(G)** The per-neuron influence of locomotion on ongoing activity minus the influence of locomotion on sound-evoked activity was significantly positive across all target-sound responsive neurons (P=3.4e-5, two-sided Wilcoxon signed-rank test) **(H)** Top: Z-scored spiking of an example neuron (black trace) overlaid on the Z-scored locomotion speed of the rat (g trace) during an example session. This neuron exhibited a correlation of R=0.44 with locomotion speed across the session. Bottom: An example from a different neuron, showing a negative correlation with locomotion speed of R=-0.4. **(I)** Distribution of spiking-locomotion speed correlation values (orange). The parallel distribution from the imaging data (Fig. 3D) is shown in light blue in the background as comparison. **(J)** Performance of stimulus detection in locomotion against state discrimination across ensembles. Blue points indicate ensembles showing significant state discrimination, red-circled points indicate ensembles showing significant stimulus detection in locomotion. Black cross shows mean±STD of the two measures. Across ensembles, both locomotive state (P=7.18e-8) and sound detection during locomotion (P=4.02e-6) could be significantly decoded from ensemble activity (Two-sided Wilcoxon signed-rank test of difference from chance of 0.5) **(K)** Discrimination of the four sound/locomotion state combinations (pre-sound in immobility, sound in immobility, pre-sound in locomotion, sound in locomotion) by ensemble activity patterns. 37/40 ensembles could significantly discriminate between these 4 options (P<0.05, bootstrap analysis)

We first examined the effect of locomotion on baseline-subtracted sound-evoked spiking responses in freely moving rats by separating responses that occurred during immobility and locomotion. While individual neurons exhibited diverse influence by locomotion, across the population of target sound-responsive neurons, sound-evoked responses were significantly weaker during locomotion as compared to immobility (**Fig. 6B,C**), consistent with our imaging data (**Fig. 2E**) and previous reports (Bigelow et al., 2019; Schneider et al., 2014).

We thus sought to test whether this locomotion-related decrease in baseline-subtracted sound-evoked responses could in part be due to increased baseline firing during locomotion as our imaging data in mouse indicated. Indeed, we found that ongoing activity, measured as the spike rate preceding stimuli presentations, was significantly higher during locomotion as compared to immobility across sound-responsive neurons (**Fig. 6D-E**). Moreover, as with our imaging data, the influence of locomotion on spiking activity in the pre-stimulus and stimulus windows were highly correlated (**Fig. 6F**, R=0.89), and significantly higher for the pre-stimulus window (**Fig. 6G**). These data suggest that increased ongoing activity during locomotion contributes to measurements of weaker sound-evoked responses in the freely-moving rat as well.

To test whether increased ongoing activity during locomotion encodes information about locomotion speed in the freely-moving rat, we examined correlations between continuous spiking activity and locomotion speed. We found similar results to the head-fixed mouse data, with the spiking activity of some neurons reliably tracking locomotion speed (**Fig. 6H**). Across the population, the distribution of correlation between neural activity and locomotion speed was skewed to the right, and highly similar to the distribution of the head-fixed data (**Fig. 6I**, skewness=0.64). These data suggest that auditory cortical neurons integrate information about locomotion speed with sound encoding during movement in the freely-moving rat. Finally, we carried out similar decoding analyses to the ones we implemented on the imaging data, to quantify the ability of neural ensembles (defined here as all simultaneously recorded putative excitatory neurons) to both discriminate locomotive state and detect sounds during locomotion. Despite having a substantially lower number of simultaneously recorded neurons as compared to the imaging data (mean±sem electro: 4.5±0.39, imaging: 29.3±4.6), we found that across ensembles, both locomotive state and sound occurrence during locomotion could be significantly decoded (**Fig. 6J**). Lastly, 37/40 ensembles could significantly discriminate between the four combinations of locomotion state and sound occurrence (**Fig. 6K**). These findings suggest that integration of sound and locomotion information by auditory cortical ensemble activity patterns generalizes across species, recording techniques and behavioral conditions.

## Discussion

In this study, we tested the hypothesis that rather than being simply suppressed during locomotion, the AC performs critical computations for sound perception in this state. In support of this hypothesis, we found that AC activity is required for sound-guided behavior during locomotion, even more than in immobility. Our neural recording experiments in both head-fixed mice and freely-moving rats revealed that underlying a net inhibitory effect of locomotion, neuronal ensembles actively and robustly encode locomotion itself in addition to sound, resulting in an integrated sound-in-motion signal.

Previous studies have found that sound-evoked responses are on average weaker during locomotion as compared to immobility, a finding we have replicated here in both head-fixed mice and freely-moving rats. A key proposed explanation for this finding is that during locomotion neural computational resources shift from auditory to visual processing (Schneider et al., 2014; Zhou et al., 2014). According to this explanation, weaker AC responses during locomotion reflect a reduced involvement of AC in sound processing in this state, in parallel to an enhancement of visual processing supported by increased responses in the visual cortex (Dadarlat and Stryker, 2017; Niell and Stryker, 2010; Vinck et al., 2015). However, the evolutionary and functional logic of this finding remains debated given the central role of sound processing during locomotion in everyday life in humans and other animals. Whether the involvement of the AC in sound processing is indeed reduced during locomotion has previously not been directly tested. Our finding that AC inactivation significantly impaired sound-guided behavior during locomotion and that this impairment was significantly larger than in immobility suggest that the AC plays an important role in sound processing during locomotion. Thus, we suspected that weaker average sound-evoked responses during locomotion reflect a different neural computation rather than a loss of function.

A hint as to the nature of the computation that AC neural ensembles perform during locomotion comes from parallel studies in other cortical regions. Specifically, although V1 responses are on average enhanced during locomotion, a number of studies have found that the influence of locomotion on visual cortical processing is better explained by sensory-motor integration than a general increase in gain. For example, one study found that locomotion modulates visual spatial integration by preferentially enhancing responses to larger visual objects (Ayaz et al., 2013). An additional study found that V1 neurons are tuned to weighted combinations of locomotion speed and the speed of the incoming visual stimulus, giving rise to multimodal locomotion-visual representations in V1 (Saleem et al., 2013). Based on these and additional studies (Fiser et al., 2016; Keller et al., 2012; Saleem et al., 2018), it has been suggested that beyond simple modulation of response magnitude, a key function of V1 is to integrate visual and locomotion information in ways that inform action and navigation (Parker et al., 2020).

Our findings suggest that cortical processing of sounds also reflect sensory-motor integration rather than simple inhibition. A first support of this possibility comes from the degree of heterogeneity in locomotion-induced modulation of activity across neurons. While early studies suggested a uniform suppression of sound-evoked responses across neurons, attributed to inhibition from secondary motor cortex (Schneider et al., 2014) and/or recruitment of local interneurons (Zhou et al., 2014), later studies observed a more heterogeneous pattern (Bigelow et al., 2019). We confirmed and extended this finding by identifying spatially intermingled subnetworks of neurons that are differentially modulated by locomotion. These data are consistent with the patterns of local heterogeneity of tone-evoked responses in the AC (Bandyopadhyay et al., 2010; Bathellier et al., 2012; Rothschild and Mizrahi, 2015; Rothschild et al., 2010; Vasquez-Lopez et al., 2017; Winkowski and Kanold, 2013), and suggest that locomotion has distinct effects on different auditory cortical subpopulations rather than inducing global suppression.

As a further test for whether AC ensembles integrate sound and locomotion, we measured whether AC neurons encode locomotion itself, in addition to sounds. When examining sound-evoked responses we observed that many neurons, as well as the average response across all responsive neurons, exhibited elevated pre-stimulus, baseline activity during locomotion. While ongoing, pre-stimulus activity is traditionally subtracted out in the calculation of sound-evoked responses (and its elevation during locomotion therefore contributes to a reduced baseline-subtracted response), we found that it conveys a highly informative signal regarding the animal’s locomotion speed. Indeed, locomotion speed coding was found to be a dominant feature of auditory cortical processing: significant encoding of locomotion speed was found in the majority of neurons and ensembles, and the combined activity of neuronal ensembles provided high-performance predictions of locomotion speed. Put together, these data suggest that locomotion speed is explicitly and reliably encoded in ongoing activity in the auditory cortex. Using a cross-validated classification approach, we found that local ensemble activity patterns significantly predicted sound occurrence, locomotive state and their combinations, suggesting co-encoding and integration of sound and locomotion information at the AC ensemble level.

Our data on the effect of locomotion on baseline-subtracted sound-evoked responses are consistent with previous studies that reported a population-average reduction (Bigelow et al., 2019; Schneider et al., 2014). However, while previous studies have suggested that this reflects a reduction of AC involvement in sound perception in favor of increased involvement of V1 in visual processing (Schneider et al., 2014; Zhou et al., 2014), our data suggests that a key underlying process is the explicit encoding and active integration of locomotion information into the sound-coding signal. Thus, our data suggest that cortical processing of sound during locomotion may in fact share common principles with cortical processing of visual and somatosensory information, in which integrative sensory-motor processing have been found to be core features s (Ayaz et al., 2013; Ayaz et al., 2019; Fiser et al., 2016; Keller et al., 2012; Parker et al., 2020; Saleem et al., 2013; Saleem et al., 2018).

Our proposed role of the auditory cortex in audio-motor integration raises the question of whether this form of integration emerges first in the cortex. Audiomotor integration exists in the inferior colliculus (Yang et al., 2020), yet whether it is a result of bottom-up activity or top-down influence from the AC remains to be determined. Furthermore, the pathways and mechanisms by which audiomotor integration in AC influences motor behavior (Xiong et al., 2015a; Znamenskiy and Zador, 2013b) remain to be further explored.

## Acknowledgments

We thank Ada Eban-Rothschild, Michael Roberts, Nancy Dess, and members of the Rothschild Lab for critical comments on earlier versions of this manuscript. This work was supported by a Whitehall Foundation Research Grant #2018-08-88 (G.R)., a NARSAD Young Investigator Grant #27668 (G.R)., a Claude D. Pepper Center Grant #AG024824 (G.R.), a National Institute of Health grant 2R01MH063649 (G.R. Co-I), and a National Institute of Health training grant T32-DC000011 (C.V.)

## Author Contributions

G.R. and C.V. designed the experiments, C.V. conducted experiments, J.L. designed behavioral systems and inactivation procedures, MC.S. and A.K. conducted behavioral experiments, C.V. and G.R. analyzed the data, G.R. wrote the manuscript with input from all authors

## METHODS

All procedures followed laboratory animal care guidelines approved by the University of Michigan Institutional Animal Care and Use Committee and conformed to National Institutes of Health guidelines.

### Animals

A total of 32 male and female Thy1-GCaMP6f mice (C57BL/6J-Tg(Thy1-GCaMP6f)GP5.17Dkim/J, JAX stock No: 025393) between the ages of 12-23 weeks were used in this study (15 in the behavioral experiments and 17 in the two-photon experiments). Mice were kept on a reverse light cycle and all imaging and behavioral sessions were performed in the dark cycle.

Data from 4 Long Evans male rats aged 4–5 months and weighing 450–550 g were also included in this study. Auditory cortical sleep data from these rats has been reported in an earlier study (Rothschild et al., 2017).

### Mouse surgery

Mice were anesthetized with Ketamine-Xylazine or isoflurane and implanted with a custom lightweight (<1 gr.) titanium head bar. For two photon calcium imaging, the muscle overlying the right auditory cortex was removed and a 3 mm diameter glass cranial window was implanted over the right auditory cortex. For the cortical inactivation experiments, small bilateral craniotomies were drilled above the auditory cortex and either 2 mm or 3 mm length custom cannulas (Plastics One, MA) were lowered into the auditory cortex. Mice received postop antibiotic ointment and Carprofen and were allowed to recover for at least 5 days before any imaging or behavioral sessions.

### Appetitive trace conditioning and AC inactivation

Mice were placed on water restriction 48 hours prior to behavioral training and received ad libitum access to food. During training and testing, mice were placed in a custom built behavioral training box, in which they were head fixed on top of a rotatable plate with an accessible water reward port. A custom Arduino-based system that received input from a rotary encoder at the base of the plate allowed presenting sounds from a speaker placed ∼10cm in front of the animal in either immobility or locomotion.

Appetitive trace conditioning in immobility: Animals were trained to associate a 1 s 8 kHz tone with subsequent water reward delivered after a delay of 1s following sound termination. Sounds (followed by water rewards) were presented following a period of continuous immobility that randomly varied across trials between 5-10 s. If the animal ran, the immobile duration counter was reset. Animals advanced to the testing phase only after they displayed consistent post-sound reward-predictive licking in locomotion for 2 consecutive days. Animals were tested following bilateral infusion of 750 nL PBS solution into AC. 24 hours following PBS infusions animals were tested following bilateral infusions of 750 nL muscimol (1 µg/µl). Unrewarded catch trials (10% of trials) were used to validate sound-triggered licking.

Appetitive trace conditioning in locomotion: A different group of mice were trained on a similar task in which sounds were presented during locomotion. Specifically, mice were trained on a task in which sounds (followed by water reward) were presented exclusively in locomotion after the animal had run a distance that randomly varied across trials between 25 and 55 20ths of a full rotation. If the animal paused for longer than 2 s then the trial was reset. Animals advanced to the testing phase only after they displayed consistent post-sound reward-predictive licking in locomotion for 2 consecutive days. Unrewarded catch trials (10% of trials) and immobility trials were used to validate sound-triggered and locomotion-specific licking, respectively. Similar to the immobility conditions, mice were first tested following infusion of 750 nL PBS solution into auditory cortex and 24 hours later following muscimol infusion.

To quantify the association between sound and subsequent water reward, we quantified the degree of increased licking in the 1 s window following sound termination and before reward delivery (0-1000 ms from sound offset) relative to the pre-sound baseline lick rate ((−1500) – (- 500) ms from sound onset). To this end we defined a “predictive lick index” as the across-trials average difference between the number of licks in the post-sound window and that of the pre-sound window.

### Two-photon imaging

During imaging sessions, mice were placed on a rotating plate while being head fixed under the microscope objective. Imaging was carried out while the head of the animal was straight, with the objective tilted using an orbital nosepiece to allow optical access to the auditory cortex. Mice were allowed to initiate movement at their leisure. Imaging was performed using an Ultima IV two-photon microscope (Bruker), a pulsed tunable laser (MaiTai eHP DeepSee by Spectra Physics) providing excitation light at 940nm and 16X or 40X water-immersion objectives (Nikon). Images (256×256 pixels) were acquired using galvanometric mirrors at ∼3 Hz to optimize signal quality and cell separation. The microscope was placed in an enclosed chamber in a dark, quiet room. Neurons were imaged at depths of 150-350 µM, corresponding to cortical L2/3.

During imaging sessions, the mouse’s behavior was video recorded using an infrared camera, which was synchronized offline with the imaging data acquisition. Locomotion and immobility were determined offline using semi-automatic movement-detection MATLAB code with manual thresholding and supervision. In addition, in most imaging sessions a rotary encoder was positioned at the base of the rotating plate allowing to acquire continuous locomotion speed. In a given daily imaging session, responses of the same neurons were imaged to multiple sound protocols. Different neuronal ensembles in the same mice were typically imaged on separate days.

Auditory stimuli were delivered via an open-field magnetic speaker (MF1, Tucker Davis Technologies) at 75 dB. The broadband noise bursts protocol consisted of 45 repeats of 1 s white noise bursts with an interstimulus interval of 3 ± 1s. The sound-masking sessions consisted of continuous presentation of broad band noise at 80 dB. The pure tone protocol (Suppl. Fig. 1) consisted of three randomly shuffled pure tones (2 kHz, 4 kHz, 8 kHz), of 20 repeats, with a duration of 1s and an interstimulus interval of 3±1s. The complex sound protocol (Suppl. Fig. 1) consisted of four randomly shuffled complex sounds (cricket, sparrow, scratch, water), with 20 or 9 repeats per stimulus. Complex stimuli duration ranged from 0.2-0.5s, padded with 0.8-0.2s of silence to create 1s long stimuli frames and an insterstimulus interval of 3±1s.

### Imaging data preprocessing and analysis

Daily imaging data of the same ensemble across multiple sound protocols was concatenated and then preprocessed using the open source Suite2P software (Pachitariu et al., 2017) for movement correction and neuronal ROI detection within the ensemble. Neural data, sound stimuli and locomotion speed signals were aligned.

Data analysis was performed using custom software written in Matlab (MathWorks).

Relative change in fluorescence (ΔF/F) across time (t) was calculated for each detected cell as 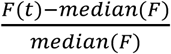, where F(t) is the mean brightness of the cell’s pixels at time t.

For determination of BBN-responsiveness of individual neurons and quantification of activity in the pre-stimulus time window (Ongoing) and stimulus time window (Evoked), the mean ΔF/F was taken across 1-4 samples preceding stimulus onset (corresponding to ∼-1.2 – 0 s), and 1-4 samples following stimulus onset (corresponding to ∼0 – 1.2 s), respectively. A cell was determined as BBN-responsive if ΔF/F during the stimulus time window was significantly higher than during the pre-stimulus time window using a one-tailed paired t-test at P<0.05 across all immobile trials. Ongoing activity levels in immobility/locomotion in the presence of background masking noise was quantified as the average ΔF/F across all time points of immobility/locomotion in the session.

A difference in BBN-evoked response magnitude between immobility and locomotion was determined using an unpaired two-sided t-test (at P<0.05) of the response magnitudes during the immobility and locomotion trials. Neurons with 8 or fewer responses in either state (immobility/locomotion) were excluded from immobility/locomotion comparisons. To determine a difference in the influence of locomotion on responses to tones and complex sounds, locomotion and immobility trials of each stimulus were compared separately.

Noise correlations between pairs of simultaneously imaged neurons were calculated as the Pearson correlation between their trial-by-trial baseline-subtracted sound responses. Cross-correlations between pairs of simultaneously imaged neurons were calculated as the cross-correlation between their continuous ΔF/F traces. Cross-correlations were normalized such that the autocorrelations at zero lag equal 1. Noise correlations and cross-correlations were calculated separately between pairs of neurons whose sound-evoked responses were (1) Both significantly enhanced during locomotion, (2) Both significantly suppressed during locomotion and (3) One neuron significantly enhanced and the other significantly suppressed during locomotion. A difference in the peak of cross-correlations between groups was tested by taking the maximum values of each cross-correlogram within a lag of ±0.66 and comparing these values across groups using a one-way ANOVA.

For calculating ΔF/F-locomotion speed correlations and speed prediction, the daily locomotion speed was smoothed using a 6-sample (∼2 s) moving average filter. The ΔF/F-locomotion speed correlation was calculated as the Pearson correlation between the continuous ΔF/F trace of each neuron and the animal’s locomotion speed.

Speed prediction was carried out using cross-validated generalized linear models on the day’s ensemble continuous activity patterns and locomotion speed. For a given ensemble, the data included the daily continuous locomotion speed and ΔF/F traces of all cells. ΔF/F traces of each cell were smoothed using a 3-sample (∼1 s) moving average filter. Locomotion speed in cm/s was log-transformed using log(*speed* + 1). In the model training phase, a random half of the daily sample points (“training set”) of locomotion and corresponding ensemble ΔF/F values were used to train a generalized linear model. In the test phase, the model used the remaining ensemble ΔF/F values (“test set”) to predict the corresponding (log) locomotion speeds. Prediction performance was quantified by the Spearman correlation between the predicted speeds and the real speeds. This procedure was repeated 200 times and the correlation values averaged across repeats to yield the final prediction performance. Repeats in which the test set included fewer than 10 non-zero speed values were excluded.

Stimulus detection during locomotion was quantified using cross-validated SVM analyses. Only ensembles with more than 10 cells and 12 trials in both immobility and locomotion were included in the decoding analyses. Data consisted of all ensemble activity patterns before BBN presentation (i.e., across-trials ensemble activity in the pre-stimulus time windows) and during BBN presentation (i.e., across-trials ensemble activity in the stimulus time windows) that occurred during locomotion. An SVM model was constructed on this data and a 10-fold cross validation was used to estimate the ability of the ensemble to discriminate between the pre-stimulus and stimulus ensemble activity patterns. Detection performance was defined as the across-fold average of percent correct predictions. To estimate significance of prediction, this procedure was performed 200 times on shuffled data identity and significant detection was determined if detection performance was higher than 95% of shuffles. Discrimination of locomotion state (immobility/locomotion) was carried out in a similar manner, but using (1) The ensemble activity patterns during the pre-stimulus time windows in immobility and (2) The pre-stimulus time windows in locomotion, as the data to be discriminated. Discrimination between the four combinations of sound occurrence and locomotion state was carried out similarly using a linear discriminant analysis, but using the ensemble activity patterns during (1) Pre-stimulus time windows in immobility (2) Pre-stimulus time windows in locomotion (3) Stimulus time windows in immobility (4) Stimulus time windows in locomotion, as the data to be discriminated. The same number of trials was included in the three models (stimulus detection during locomotion, state discrimination and stimulus+state discrimination) by removing excess trials in the data with more trials.

Analysis of the electrophysiology data was carried out similarly to the imaging data, but using spike counts instead of ΔF/F as the neural measure and using a stimulus response time window of 1-450 ms and a pre-stimulus time window of -450-0 ms relative to sound onset. Neurons with 10 or fewer responses in either state (immobility/locomotion) were excluded from immobility/locomotion comparisons. A minimum of 20 trials in both immobility and locomotion were required for inclusion in the decoding analyses. Spiking-speed correlations were calculated by binning spiking and speeds into 200 ms bins and calculating the Spearman correlation. Discrimination between the 4 state/sound combinations were carried out using a pseudolinear discriminant analysis.

### Rat pretraining, surgery and electrophysiological recordings

The rat behavioral and surgery procedures have been described previously (Znamenskiy and Zador, 2013a). Briefly, after habituation to daily handling over several weeks, rats were pretrained to run on an E-shaped raised track for liquid food rewards (sweetened condensed milk). Rats were then implanted with a microdrive array with 21 independently moveable tetrodes (groups of four twisted 12.5 μm nichrome wires assembled in a bundle). Seven tetrodes were targeted to the left primary AC (−4.8 mm AP, 5.5 mm ML, 25° lateral from midline). Other tetrodes targeted left dorsal CA1 region of the hippocampus and left PFC, but these data are not included here. Over the course of two weeks following implantation, AC tetrodes were advanced gradually and responses to sound stimuli were used to validate approach to primary AC.

Data were collected using the NSpike data acquisition system (L.M. Frank and J. MacArthur, Harvard Instrumentation Design Laboratory). Spike data were sampled at 30 kHz, digitally filtered between 300 Hz and 6 kHz (two-pole Bessel for high and low pass) and threshold crossing events were saved to disk (40 samples at 30 kHz). Individual units (putative single neurons) were identified by clustering spikes using peak amplitude, principal components and spike width as variables (MatClust). Behavior sessions were recorded with an overhead monochrome CCD camera (30 fps) and the animal’s position and speed were detected using an infrared light emitting diode array with a large and a small cluster of diodes attached to the preamps. For binary assignment of immobility and locomotion we used a standard 4 cm/s speed threshold.

Approximately 14 d after implantation, animals were introduced to the Y-track and data gathering commenced. Animals were trained on the Y-track for 10–12 d in 3–4 20-min training sessions per day with interleaving 20-to 30-min sleep sessions in the rest box. Data for each neuron was pooled across daily sessions. During training sessions, sweetened condensed milk rewards were automatically delivered in food wells triggered by animal’s nose-poke crossing of an IR beam. Rats initiated each trial by a nose-poke in the home well and receiving a reward. In ∼75% of trials the next reward was delivered in the silent well if the rat nose-poked there. In a pseudorandom ∼25% of trials (sound trials separated by 2–5 silent trials), 5 s after nose-poking in the home arm, a target sound series was emitted from a speaker, indicating that the next reward would be delivered in the sound well if the rat next nose-poked there. The speaker was placed at the end of the sound arm in the first days of training and moved to the center junction after rats displayed consistent correct choices in more than ∼70% of trials. The target sound was a pair of upward chirps, consisting of one 200-ms chirp with frequency modulated from 3 to 4 kHz, an interchirp interval of 50 ms, and a second 200-ms chirp with frequency modulated from 9 to 12 kHz. The series of target sounds was presented at 1 Hz and stopped after 12 s or once the rat made a correct or incorrect choice by a nose-poke in one of the wells. Reward amount in the sound well was double the reward amount in the home or silent well. Following the Y-track training days, two rats continued to perform the same task on a W-shaped track for an additional 3 days.

## Data and code availability

Source data and analysis scripts have been deposited with FigShare (10.6084/m9.figshare.19750678).

---

**Supplementary Fig. 1:**
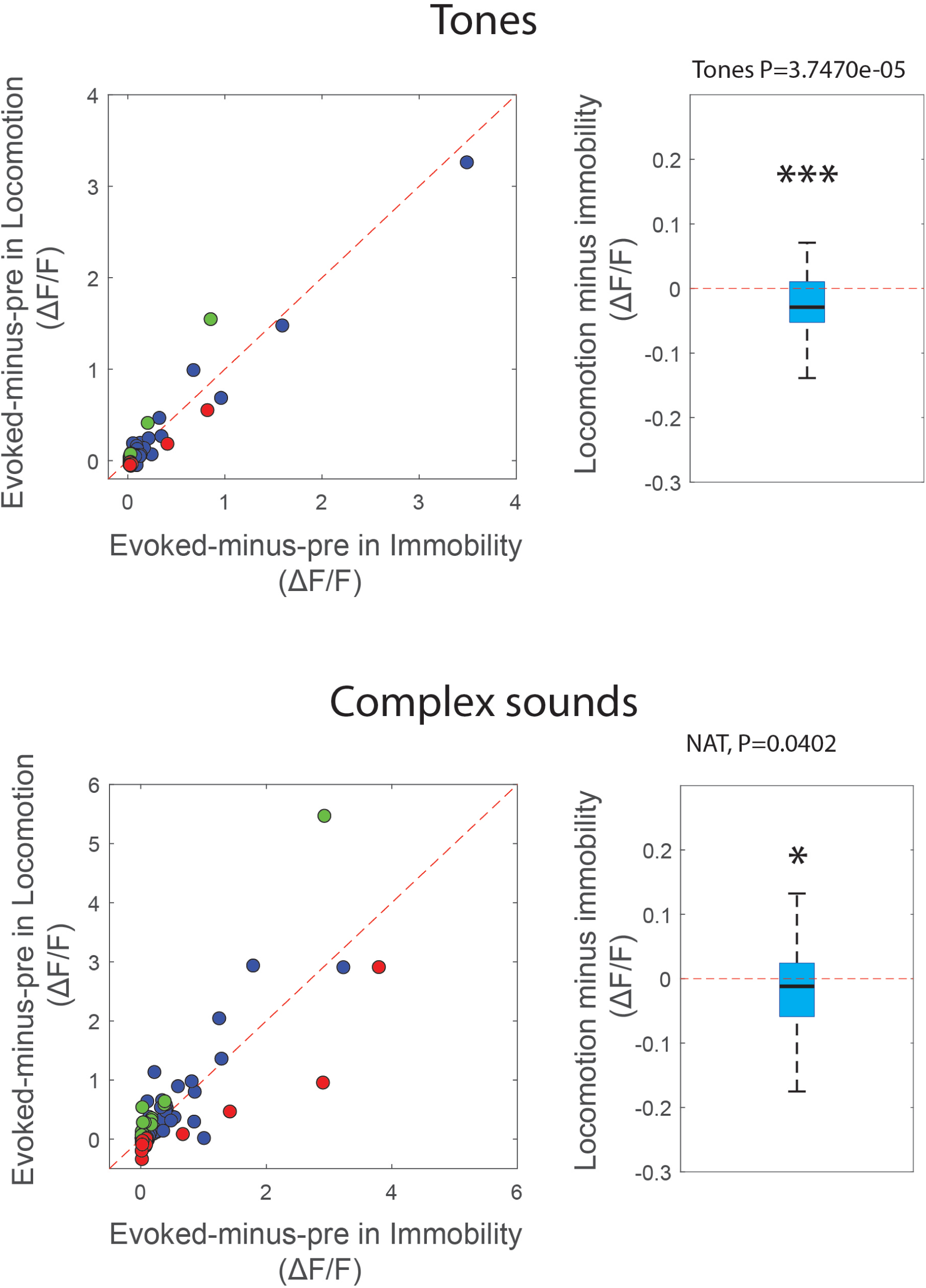
Locomotion influence on AC responses to tones and complex sounds. Baseline-subtracted sound-evoked responses in immobility and locomotion for tones (top) and complex sounds (bottom). Graphical conventions same as Fig. 1E. While individual responses showed diversity in locomotion-related influence, population-level responses to both tones and complex sounds were significantly reduced during locomotion (Tones: P=3.7e-5, Complex sounds P=0.0402, two-sided Wilcoxon signed-rank test).

**Supplementary Fig. 2:**
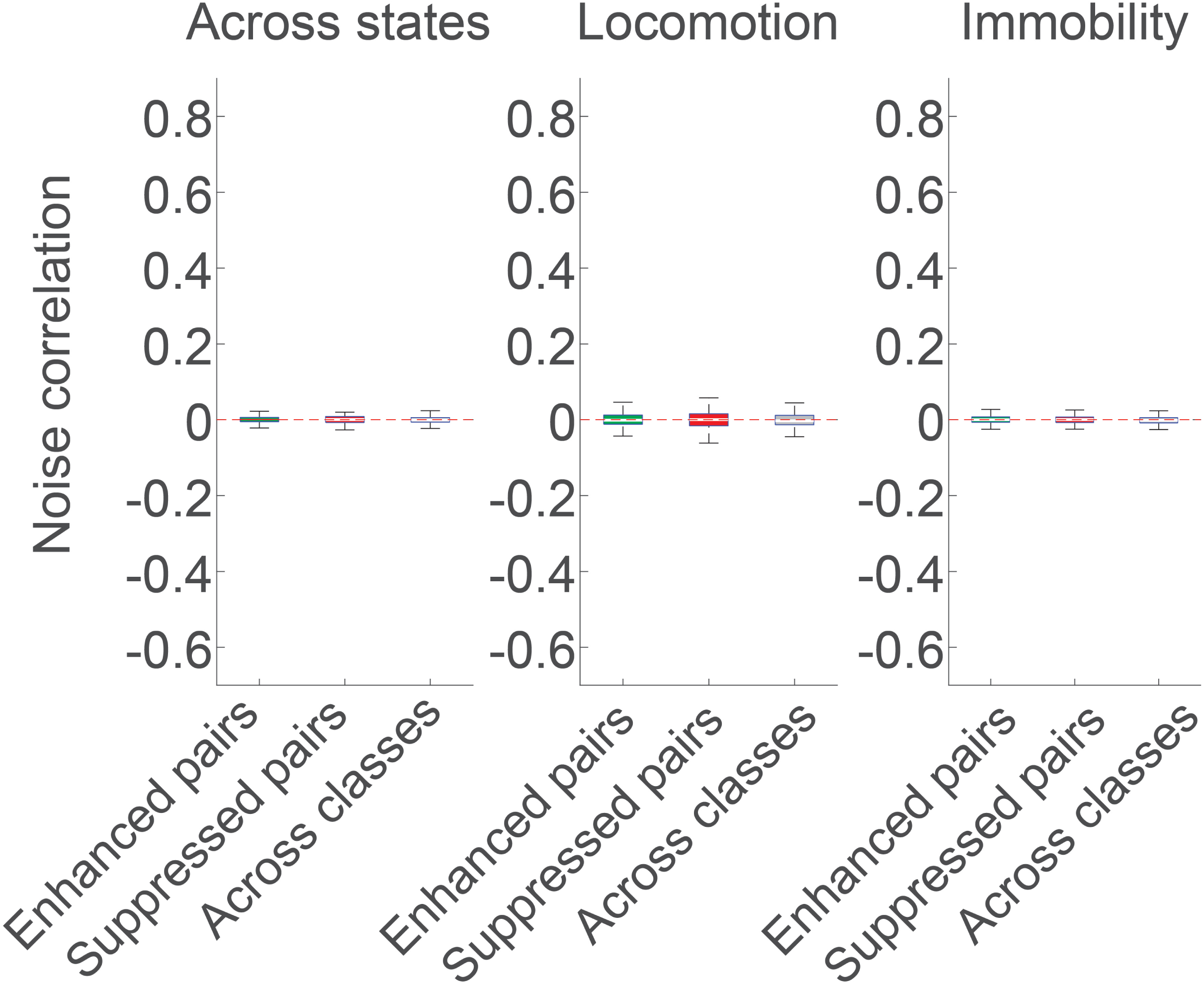
Noise correlations of trial-shuffled data. Data parallels Fig. 2G but following trial shuffling. No significant differences were observed.

**Supplementary Fig. 3:**
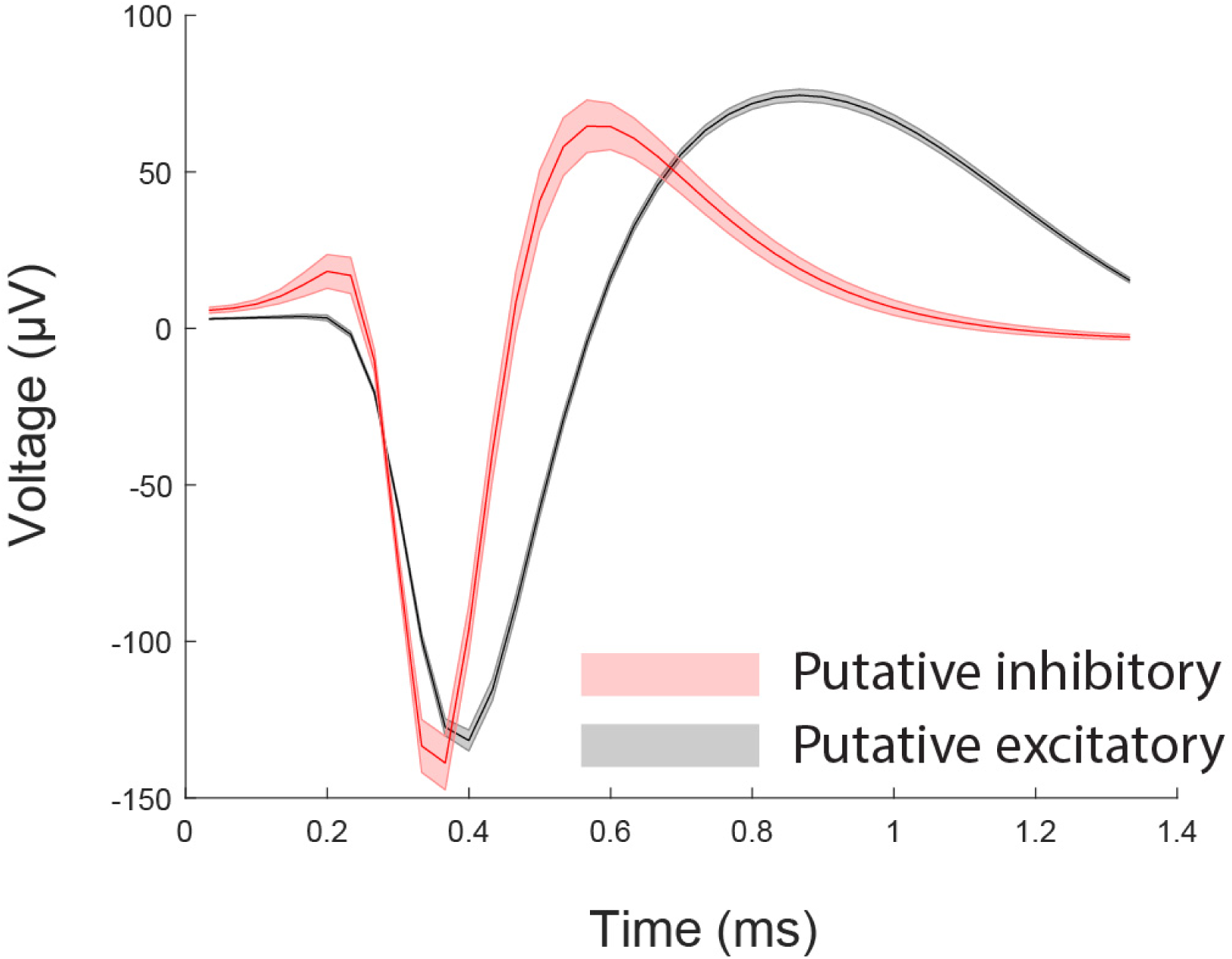
Spike waveforms of putative excitatory and inhibitory neurons. Traces show mean±SEM of spikes from the corresponding classes. Putative inhibitory fast-spiking interneurons were excluded from analyses.

## References

1. Ayaz, A., Saleem, A.B., Scholvinck, M.L., and Carandini, M. (2013). Locomotion controls spatial integration in mouse visual cortex. Curr Biol 23, 890–894.

2. Ayaz, A., Stauble, A., Hamada, M., Wulf, M.A., Saleem, A.B., and Helmchen, F. (2019). Layer-specific integration of locomotion and sensory information in mouse barrel cortex. Nat Commun 10, 2585.

3. Bandyopadhyay, S., Shamma, S.A., and Kanold, P.O. (2010). Dichotomy of functional organization in the mouse auditory cortex. Nat Neurosci 13, 361–368.

4. Bathellier, B., Ushakova, L., and Rumpel, S. (2012). Discrete neocortical dynamics predict behavioral categorization of sounds. Neuron 76, 435–449.

5. Bigelow, J., Morrill, R.J., Dekloe, J., and Hasenstaub, A.R. (2019). Movement and VIP Interneuron Activation Differentially Modulate Encoding in Mouse Auditory Cortex. eNeuro 6.

6. Carr, K.W., Tierney, A., White-Schwoch, T., and Kraus, N. (2016). Intertrial auditory neural stability supports beat synchronization in preschoolers. Dev Cogn Neuros-Neth 17, 76–82.

7. Chen, J.L., Penhune, V.B., and Zatorrel, R.J. (2008). Moving on time: Brain network for auditory-motor synchronization is modulated by rhythm complexity and musical training. J Cognitive Neurosci 20, 226–239.

8. Cohen, L., Rothschild, G., and Mizrahi, A. (2011). Multisensory integration of natural odors and sounds in the auditory cortex. Neuron 72, 357–369.

9. Cornwell, T., Woodward, J., Wu, M.M., Jackson, B., Souza, P., Siegel, J., Dhar, S., and Gordon, K.E. (2020). Walking With Ears: Altered Auditory Feedback Impacts Gait Step Length in Older Adults. Frontiers in Sports and Active Living 2.

10. Cuppone, A.V., Cappagli, G., and Gori, M. (2018). Audio Feedback Associated With Body Movement Enhances Audio and Somatosensory Spatial Representation. Frontiers in Integrative Neuroscience 12.

11. Dadarlat, M.C., and Stryker, M.P. (2017). Locomotion Enhances Neural Encoding of Visual Stimuli in Mouse V1. J Neurosci 37, 3764–3775.

12. Falk, B., Jakobsen, L., Surlykke, A., and Moss, C.F. (2014). Bats coordinate sonar and flight behavior as they forage in open and cluttered environments. J Exp Biol 217, 4356–4364.

13. Fiser, A., Mahringer, D., Oyibo, H.K., Petersen, A.V., Leinweber, M., and Keller, G.B. (2016). Experience-dependent spatial expectations in mouse visual cortex. Nat Neurosci 19, 1658–1664.

14. Fox, M.W. (1984). The whistling hunters : field studies of the Asiatic wild dog (Cuon alpinus) (Albany: State University of New York Press).

15. Ghose, K., Horiuchi, T.K., Krishnaprasad, P.S., and Moss, C.F. (2006). Echolocating bats use a nearly time-optimal strategy to intercept prey. Plos Biol 4, 865–873.

16. Jaramillo, S., and Zador, A.M. (2011). The auditory cortex mediates the perceptual effects of acoustic temporal expectation. Nat Neurosci 14, 246–U340.

17. Karpati, F.J., Giacosa, C., Foster, N.E.V., Penhune, V.B., and Hyde, K.L. (2015). Dance and the brain: a review. Ann Ny Acad Sci 1337, 140–146.

18. Keller, G.B., Bonhoeffer, T., and Hubener, M. (2012). Sensorimotor mismatch signals in primary visual cortex of the behaving mouse. Neuron 74, 809–815.

19. Kuchibhotla, K.V., Gill, J.V., Lindsay, G.W., Papadoyannis, E.S., Field, R.E., Sten, T.A.H., Miller, K.D., and Froemke, R.C. (2017). Parallel processing by cortical inhibition enables context-dependent behavior. Nat Neurosci 20, 62–71.

20. Moss, C.F., and Surlykke, A. (2001). Auditory scene analysis by echolocation in bats. J Acoust Soc Am 110, 2207–2226.

21. Niell, C.M., and Stryker, M.P. (2010). Modulation of visual responses by behavioral state in mouse visual cortex. Neuron 65, 472–479.

22. Pachitariu, M., Stringer, C., Dipoppa, M., Schröder, S., Rossi, L.F., Dalgleish, H., Carandini, M., and Harris, K.D. (2017). Suite2p: beyond 10,000 neurons with standard two-photon microscopy. bioRxiv, 061507.

23. Parker, P.R.L., Brown, M.A., Smear, M.C., and Niell, C.M. (2020). Movement-Related Signals in Sensory Areas: Roles in Natural Behavior. Trends Neurosci 43, 581–595.

24. Ravignani, A., and Cooke, P.F. (2016). The evolutionary biology of dance without frills. Current Biology 26, R878–R879.

25. Redd, C.B., and Bamberg, S.J.M. (2012). A Wireless Sensory Feedback Device for Real-Time Gait Feedback and Training. Ieee-Asme T Mech 17, 425–433.

26. Rodger, M.W.M., Young, W.R., and Craig, C.M. (2014). Synthesis of Walking Sounds for Alleviating Gait Disturbances in Parkinson’s Disease. Ieee T Neur Sys Reh 22, 543–548.

27. Rodgers, C.C., and DeWeese, M.R. (2014). Neural Correlates of Task Switching in Prefrontal Cortex and Primary Auditory Cortex in a Novel Stimulus Selection Task for Rodents. Neuron 82, 1157–1170.

28. Rothschild, G., Eban, E., and Frank, L.M. (2017). A cortical-hippocampal-cortical loop of information processing during memory consolidation. Nat Neurosci 20, 251–259.

29. Rothschild, G., and Mizrahi, A. (2015). Global order and local disorder in brain maps. Annu Rev Neurosci 38, 247–268.

30. Rothschild, G., Nelken, I., and Mizrahi, A. (2010). Functional organization and population dynamics in the mouse primary auditory cortex. Nat Neurosci 13, 353–360.

31. Saleem, A.B., Ayaz, A., Jeffery, K.J., Harris, K.D., and Carandini, M. (2013). Integration of visual motion and locomotion in mouse visual cortex. Nat Neurosci 16, 1864–1869.

32. Saleem, A.B., Diamanti, E.M., Fournier, J., Harris, K.D., and Carandini, M. (2018). Coherent encoding of subjective spatial position in visual cortex and hippocampus. Nature 562, 124–127.

33. Schauer, M., and Mauritz, K.-H. (2003). Musical motor feedback (MMF) in walking hemiparetic stroke patients: randomized trials of gait improvement. Clinical Rehabilitation 17, 713–722.

34. Schneider, D.M., Nelson, A., and Mooney, R. (2014). A synaptic and circuit basis for corollary discharge in the auditory cortex. Nature 513, 189–194.

35. Tajadura-Jiménez, A., Basia, M., Deroy, O., Fairhurst, M., Marquardt, N., and Bianchi-Berthouze, N. (2015). As Light as your Footsteps: Altering Walking Sounds to Change Perceived Body Weight, Emotional State and Gait. CHI ‘15 Proceedings of the 33rd Annual ACM Conference on Human Factors in Computing Systems, 2943–2952.

36. Tierney, A., and Kraus, N. (2013). The Ability to Move to a Beat Is Linked to the Consistency of Neural Responses to Sound. Journal of Neuroscience 33, 14981–14988.

37. Tierney, A., and Kraus, N. (2016). Getting back on the beat: links between auditory-motor integration and precise auditory processing at fast time scales. Eur J Neurosci 43, 782–791.

38. Triblehorn, J.D., and Yager, D.D. (2005). Timing of praying mantis evasive responses during simulated bat attack sequences. J Exp Biol 208, 1867–1876.

39. Turchet, L., Camponogara, I., and Cesari, P. (2015). Interactive footstep sounds modulate the perceptual-motor aftereffect of treadmill walking. Exp Brain Res 233, 205–214.

40. Turchet, L., Camponogara, I., Nardello, F., Zamparo, P., and Cesari, P. (2018). Interactive footsteps sounds modulate the sense of effort without affecting the kinematics and metabolic parameters during treadmill-walking. Appl Acoust 129, 379–385.

41. Turchet, L., Serafin, S., and Cesari, P. (2013). Walking Pace Affected by Interactive Sounds Simulating Stepping on Different Terrains. Acm T Appl Percept 10.

42. Ulanovsky, N., Las, L., and Nelken, I. (2003). Processing of low-probability sounds by cortical neurons. Nat Neurosci 6, 391–398.

43. Vasquez-Lopez, S.A., Weissenberger, Y., Lohse, M., Keating, P., King, A.J., and Dahmen, J.C. (2017). Thalamic input to auditory cortex is locally heterogeneous but globally tonotopic. Elife 6.

44. Vinck, M., Batista-Brito, R., Knoblich, U., and Cardin, J.A. (2015). Arousal and locomotion make distinct contributions to cortical activity patterns and visual encoding. Neuron 86, 740–754.

45. Whitton, J.P., Hancock, K.E., and Polley, D.B. (2014). Immersive audiomotor game play enhances neural and perceptual salience of weak signals in noise. Proc Natl Acad Sci U S A 111, E2606–2615.

46. Winkowski, D.E., and Kanold, P.O. (2013). Laminar transformation of frequency organization in auditory cortex. J Neurosci 33, 1498–1508.

47. Xiong, Q., Znamenskiy, P., and Zador, A.M. (2015a). Selective corticostriatal plasticity during acquisition of an auditory discrimination task. Nature 521, 348–351.

48. Xiong, Q.J., Znamenskiy, P., and Zador, A.M. (2015b). Selective corticostriatal plasticity during acquisition of an auditory discrimination task. Nature 521, 348-+.

49. Yang, Y., Lee, J., and Kim, G. (2020). Integration of locomotion and auditory signals in the mouse inferior colliculus. Elife 9.

50. Zhou, M., Liang, F., Xiong, X.R., Li, L., Li, H., Xiao, Z., Tao, H.W., and Zhang, L.I. (2014). Scaling down of balanced excitation and inhibition by active behavioral states in auditory cortex. Nat Neurosci 17, 841–850.

51. Znamenskiy, P., and Zador, A.M. (2013a). Corticostriatal neurons in auditory cortex drive decisions during auditory discrimination. Nature 497, 482-+.

52. Znamenskiy, P., and Zador, A.M. (2013b). Corticostriatal neurons in auditory cortex drive decisions during auditory discrimination. Nature 497, 482–485.

